# The genomic landscape of *Saccharomyces paradoxus* introgression in geographically diverse *Saccharomyces cerevisiae* strains

**DOI:** 10.1101/2022.08.01.502362

**Authors:** Anne Clark, Maitreya J. Dunham, Joshua M. Akey

## Abstract

Natural hybrids between many pairs of the *Saccharomyces* species have been observed. Hybridization can allow for rapid adaptation to new environments, and when followed by repeated backcrossing can lead to small pieces of introgressed sequence from an individual of one species remaining in the genome of an individual of another species. Introgressed sequences that persist over time are interesting both because they provide evidence of past hybridization events and because they may contribute to functional and phenotypic diversity. Previous studies have identified some examples of introgressed sequences in *S. cerevisiae*, but we sought to gain a more comprehensive understanding of the landscape of introgression in this species. We developed a simple, flexible approach based on a hidden Markov model to identify introgression in yeast genomes. We used our method to look for introgression in 93 diverse *S. cerevisiae* genomes from its closest relative, *S. paradoxus*. We found evidence of introgression in all strains we considered, but the amount and location of introgression varied widely. We show that introgression contributes substantially to the total genetic diversity within these strains, and has the potential to confound inferences of their evolutionary relationships. Further characterizing introgression across diverse *Saccharomyces* species may help us better understand their evolution and the role that hybridization has played in adaptation to new environments.

## Introduction

Since the original formulation of the theory of evolution, it has been apparent that the evolution of different species cannot be understood in isolation. Charles Darwin wrote of the “complex relations of all animals and plants to each other in the struggle for existence,” drawing on many examples of the evolution of one species being influenced by another (Darwin 1859). As species compete or cooperate in shared environments, their interactions influence the composition of the genomes that evolve. But species can also affect each other’s genomes more directly via hybridization or horizontal gene transfer. The ongoing improvement of sequencing technology has enabled more detailed studies of the movement of genetic material between species (Goulet et al. 2017).

The acquisition of genetic material from another species or strain can allow for more rapid adaptation to new environments by taking advantage of existing evolutionary innovation. For example, horizontally transferred genes have allowed for the evolution of multiple–drug– resistant *Staphylococcus aureus* (Price et al. 2012). In sexual organisms, hybridization can also facilitate adaptation, and hybrids are frequently useful in agricultural and industrial applications (Goulet et al. 2017; Krogerus et al. 2017; Peris et al. 2018). When hybridization is followed by repeated backcrossing, it can result in genomes that are mainly one species with interspersed introgressed sequences remaining from the other species. Introgressed sequences that originated from relatively old hybridizations but are still present in modern genomes may have remained because they are adaptive (Anderson and Stebbins 1954; Goulet et al. 2017).

The *Saccharomyces* yeast species were originally defined by the relatively low spore viability between them; this viability is not zero (Peris et al. 2018), however, and the diversity of naturally-occurring hybrids that has been observed suggests that they hybridize relatively frequently (Morales and Dujon 2012). One of the first such hybrids was found in the common brewing strain *S. carlsbergensis* (now *S. pastorianus*), and more extensive sequencing has since revealed additional hybrids within the genus, as well as a number of shorter introgressed sequences (Morales and Dujon 2012). In particular, the sequencing of a great variety of strains of multiple *Saccharomyces* species has made it possible to consider global patterns of introgression among these organisms, and to examine how this introgression may have influenced their evolution.

In addition to including the most well-characterized laboratory strain, *S. cerevisiae* has had the most strains sequenced out of all the *Saccharomyces* species. In particular, the 100-genomes strain collection provided 93 new *S. cerevisiae* genomes assembled *de novo*, with quality approaching that of the S288c reference genome (Strope et al. 2015). Hundreds of other wild, fermentation, and clinical strains have also been sequenced in the past few years (Duan et al. 2018; Zhu et al. 2016; Gallone et al. 2016; Peter et al. 2018), though these have most often been assembled using S288c as a reference.

*S. paradoxus* is on average ~14% diverged from *S. cerevisiae* at the nucleotide level. Examples of *S. paradoxus* introgressions in *S. cerevisiae* have been identified; these have mainly consisted of very large segments (Morales and Dujon 2012) or individual genes (Strope et al. 2015). These analyses suggest that introgression is widespread in *S. cerevisiae*. However, the methodological approaches used to identify introgression in yeast have relied on sequence identity—broadly, searching for regions of the genome that match a *S. paradoxus* reference better than a *S. cerevisiae* one. Although this pattern is a reasonable expectation for introgressed sequence, it may also be a consequence of incomplete lineage sorting (Goulet et al. 2017; Maddison et al. 2006). To this end, we first used coalescent theory to establish that relying on sequence identity is appropriate given the evolutionary history of these organisms.

We then developed a hidden Markov model (HMM)–based method to predict the species of origin of each site in a yeast genome. This approach allowed us to infer the boundaries of introgressed regions more precisely, instead of focusing only on individual genes. We evaluated the performance of our method on simulated sequence data, and found it to be faster, more accurate, and simpler to parameterize than an existing phylo-HMM-based method for identifying introgression.

We used our HMM-based approach to search for *S. paradoxus* introgression in the set of 93 diverse *S. cerevisiae* strains sequenced for the 100-genomes collection (Strope et al. 2015). We found some level of introgression on nearly every chromosome of every strain considered, though strains vary substantially in their amount of introgression. Overall, the introgressions we predict represent a substantial contribution to diversity, accounting for approximately 7% of the total nucleotide diversity among the strains. In some cases, introgressed sequences can influence our inference of the phylogenetic relationship of the strains. We identify genes previously suggested to be introgressed, but also identify many novel introgressions. In general, this method has allowed us to characterize the global distribution of *S. paradoxus* introgression across a set of diverse *S. cerevisiae* genomes, and may allow us to further expand our understanding of introgression within the *Saccharomyces* yeasts.

## Results

### Establishing the validity of sequence identity-based approaches for identifying introgression

Incomplete lineage sorting (ILS) can occur when lineages within a species coalesce farther back in time than when that species diverged from another species (Maddison 1997). When such ancient coalescences exist, individuals from different species may look more similar to each other than to other members of their own species (Figure 1a), potentially resulting in false positives when using sequence identity to infer the presence of introgression. But ILS is only likely to occur when the time of divergence is recent relative to the effective population sizes of the species (Wakeley 2009). Looking backward in time, when the species are more diverged there is more time for within-species coalescences to finish before the species join, so ILS is less likely (Figure 1b); conversely, when population sizes are larger the coalescences finish less quickly, so ILS is more likely. Given the divergence time and effective population sizes of two species, we can analytically calculate the probability of ILS for these species using equations based on coalescent theory (Rosenberg 2002).

**Figure 1.**
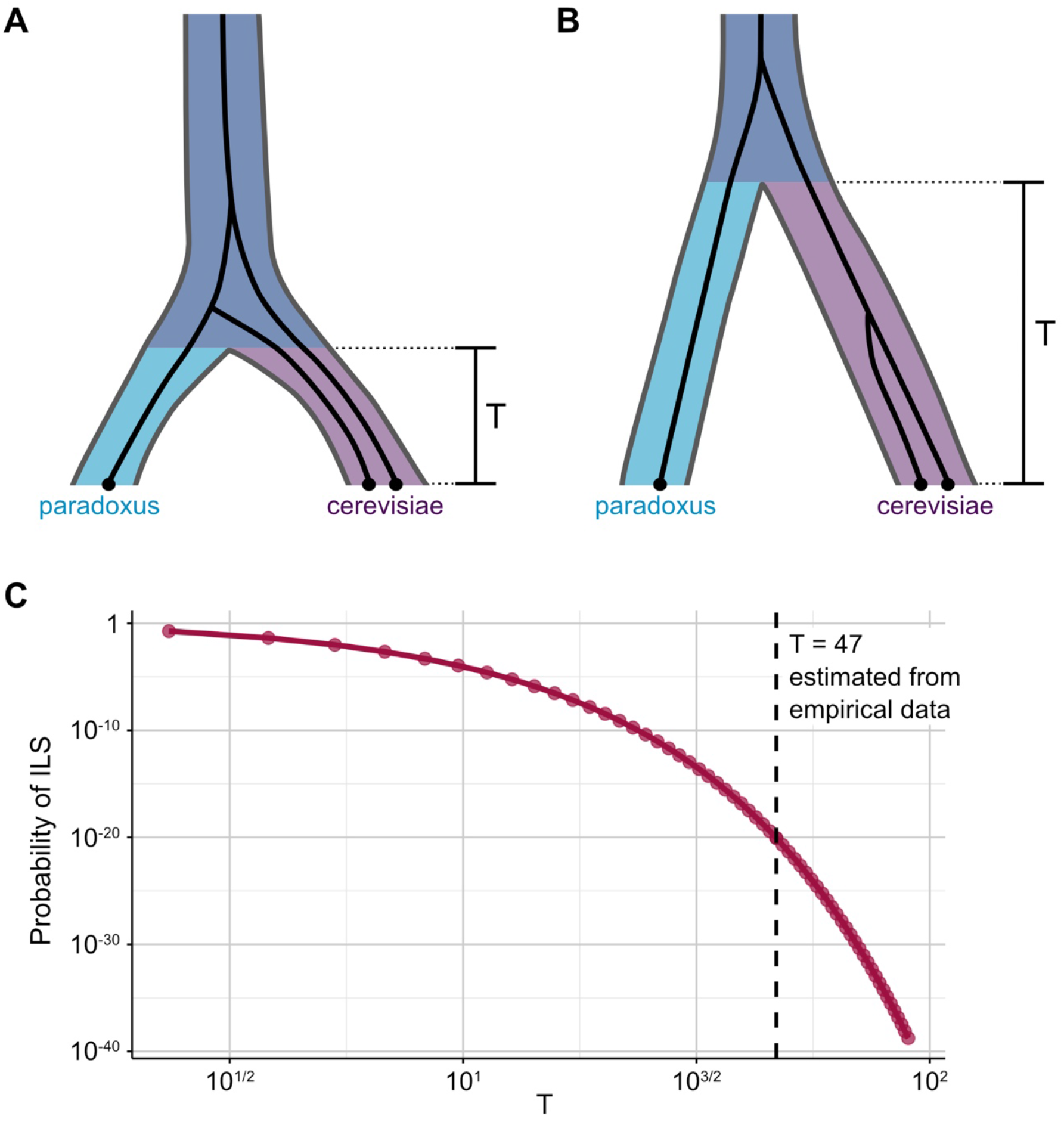
The probability of ILS in *S. cerevisiae* and *S. paradoxus*. (*A*) ILS can occur when the divergence time between species is short relative to their effective population sizes. (*B*) A longer divergence time (or a smaller population size) makes it more likely that coalescences occur within species than between them. (*C*) We calculate the probability of ILS for a range of values of the scaled divergence time—which is directly proportional to the actual divergence time, but inversely proportional to the effective population size. For our best estimate of T = 47, the probability of ILS is very small (~10^−20^), but for a value of T an order of magnitude smaller, the probability of ILS is substantial (~10^−2^).

Although the divergence time and effective population sizes for *S. cerevisiae* and *S. paradoxus* are not known with certainty, we find that the probability of ILS is essentially very small for much of the parameter space that is plausible based on empirical data (Figure 1c). For example, assuming an effective population size of 8×10^6^ for each species (Fay and Benavides 2005) and a divergence time of 3.75×10^8^ generations (see Methods), the probability of ILS is ~10^−20^. But if we allow for uncertainty in our parameter estimates by shortening the divergence time or increasing the effective population size, the probability of ILS increases. In general, however, the probability of ILS is small over the most likely range of the parameter space. Thus, these data show that ILS is unlikely to significantly confound detection of introgressed *S. paradoxus* sequences in *S. cerevisiae* and justify the use of sequence based approaches.

### Developing and evaluating a hidden Markov model to detect introgressed sequences

To detect introgression, we implemented a hidden Markov model (HMM) that has one state for each reference species (*S. cerevisiae* and *S. paradoxus*) in addition to one unknown state (Figure 1a). The model is used to assign a state to each alignment column in a given three-way chromosome alignment. Only alignment columns that are polymorphic and contain no gaps are considered.

In order to rigorously evaluate our method, we tested its performance on sequences with introgression generated using the coalescent simulator ms (Hudson 2002) with a variety of migration rates. We evaluated the true and false positives rates of our method (Figure 2b), as well as those of an existing method for detecting introgression, PhyloNet-HMM (Liu et al. 2014). Instead of simply inferring the species of origin of individual alignment columns, PhyloNet-HMM infers the underlying gene tree and species tree, which results in six possible states at each site compared to our three. As a result of this additional complexity, our method requires fewer parameter estimates and also runs ~40x faster on the simulated sequence data.

**Figure 2.**
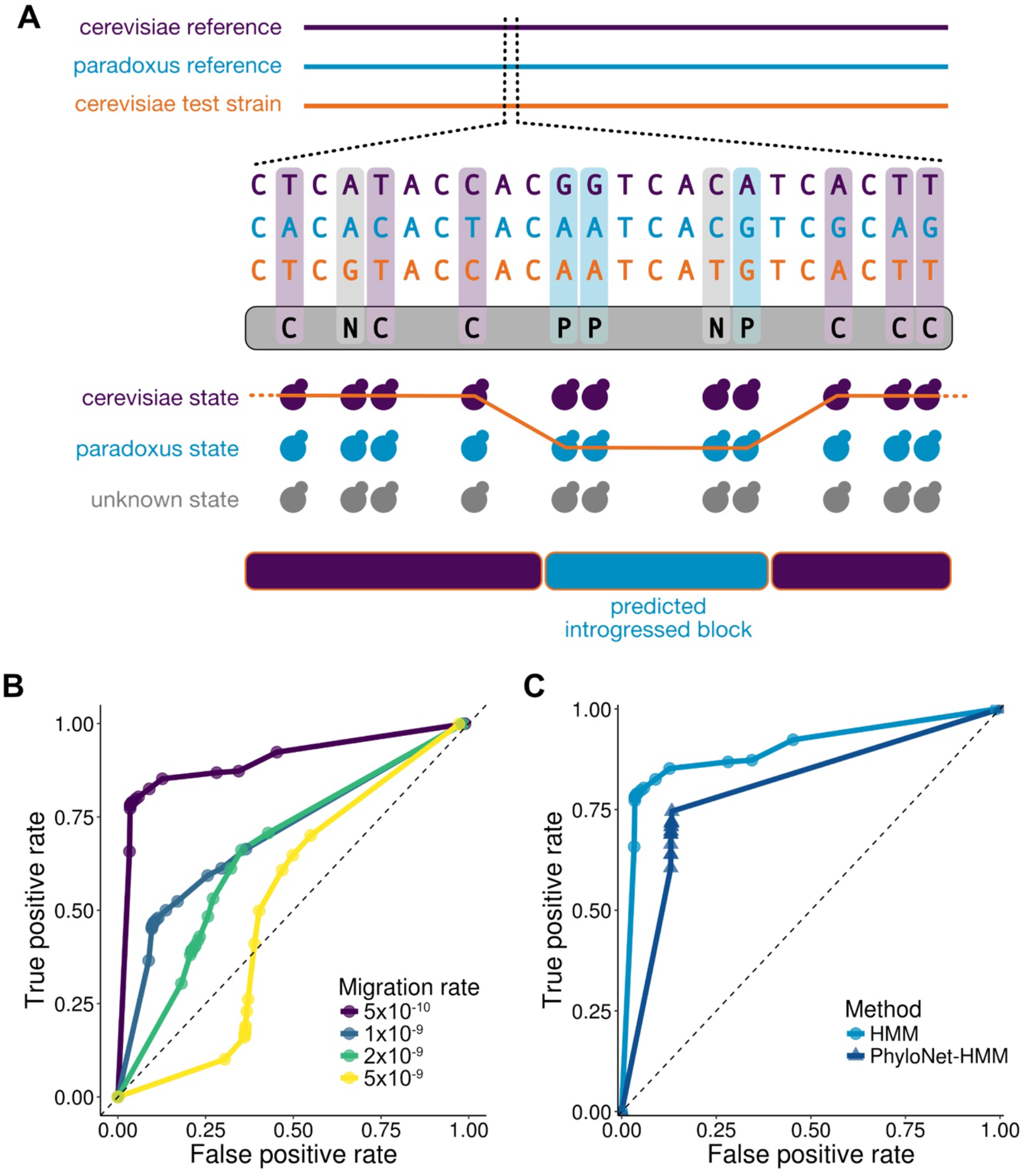
Structure and performance of our HMM for identifying introgression. (*A*) The chromosome of the current test strain is aligned to the corresponding chromosomes of the reference strains, and polymorphic alignment columns are coded by whether the test strain matches each of the references. The HMM is then used to assign a state to each of these alignment columns, with the paradoxus state indicating introgression. Consecutive occurrences of each state are grouped into regions. (*B*) True positive versus false positive rate for our method for a variety of migration rates, where the units are the fraction of the *S. cerevisiae* population made up of *S. paradoxus* individuals in each generation. Our method performs worse for higher migration rates, primarily because the high level of introgression in the reference *cerevisiae* sequence makes it difficult to identify introgression in the test sequence. (*C*) Comparison of performance of our HMM method and PhyloNet–HMM for a migration rate of 5×10^−10^. Our method outperformed PhyloNet–HMM for all migration rates tested, as shown in Supplemental Figure 1.

Both methods were run on sets of sequences generated using a range of migration rates. The performance of both is highly dependent on the specific parameters used to simulate the sequences; in particular, for higher migration rates, the level of introgression in the reference sequence makes it difficult for either method to detect introgression. For all sets of sequences analyzed, however, our method outperformed PhyloNet-HMM (Figure 2c).

### Landscape of *paradoxus*–like introgressed sequences in *cerevisiae* strains and comparison to previously–identified introgressions

We analyzed the genomes of the 93 strains that were newly sequenced for the 100-genomes collection (Strope et al. 2015). Each chromosome of each strain was aligned to *S. cerevisiae* S288c and *S. paradoxus* CBS432 using MAFFT (Katoh and Standley 2013). Using our hidden Markov model-based approach, we predict introgressions on every chromosome (Figure 2a). In total, we identify 3,147 introgressed regions across the 93 strains, ranging in size from 27 to 35,072 bp (median = 149 bp, mean = 1,071 bp; Supplemental Table 1). These introgressions collectively cover approximately 7.7% of the genome.

As expected, the introgressions we identify generally match the *S. paradoxus* reference more closely than the *S. cerevisiae* one, with the greatest concentration falling near 100% identity with *S. paradoxus* and 85-90% identity with *S. cerevisiae* (Supplemental Figure 2). However, some regions we identify do not match the *S. paradoxus* reference well. For example, approximately 30% of regions have a sequence identity of less than 90% to the *S. paradoxus* reference, but because these tend to be shorter regions, they only account for approximately 6% of introgressed bases we identify. These regions may be a better match to another *S. paradoxus* strain or another species—or may just be in poorly–aligned regions.

Previously, 287 genes were identified as putatitively introgressed from *S. paradoxus* in these 93 strains (Strope et al. 2015); these genes were found by individually comparing the sequence identity with *S. cerevisiae* and *S. paradoxus* references across the entire gene to predefined thresholds. We identify nearly all of these genes, as well as an additional 201 genes (Figure 3b). The five genes we fail to identify are due to translocations that we do not account for in our alignments, while the additional 201 genes we identify are mainly due to our ability to identify smaller parts of genes that appear to be introgressed (Figure 3c).

**Figure 3.**
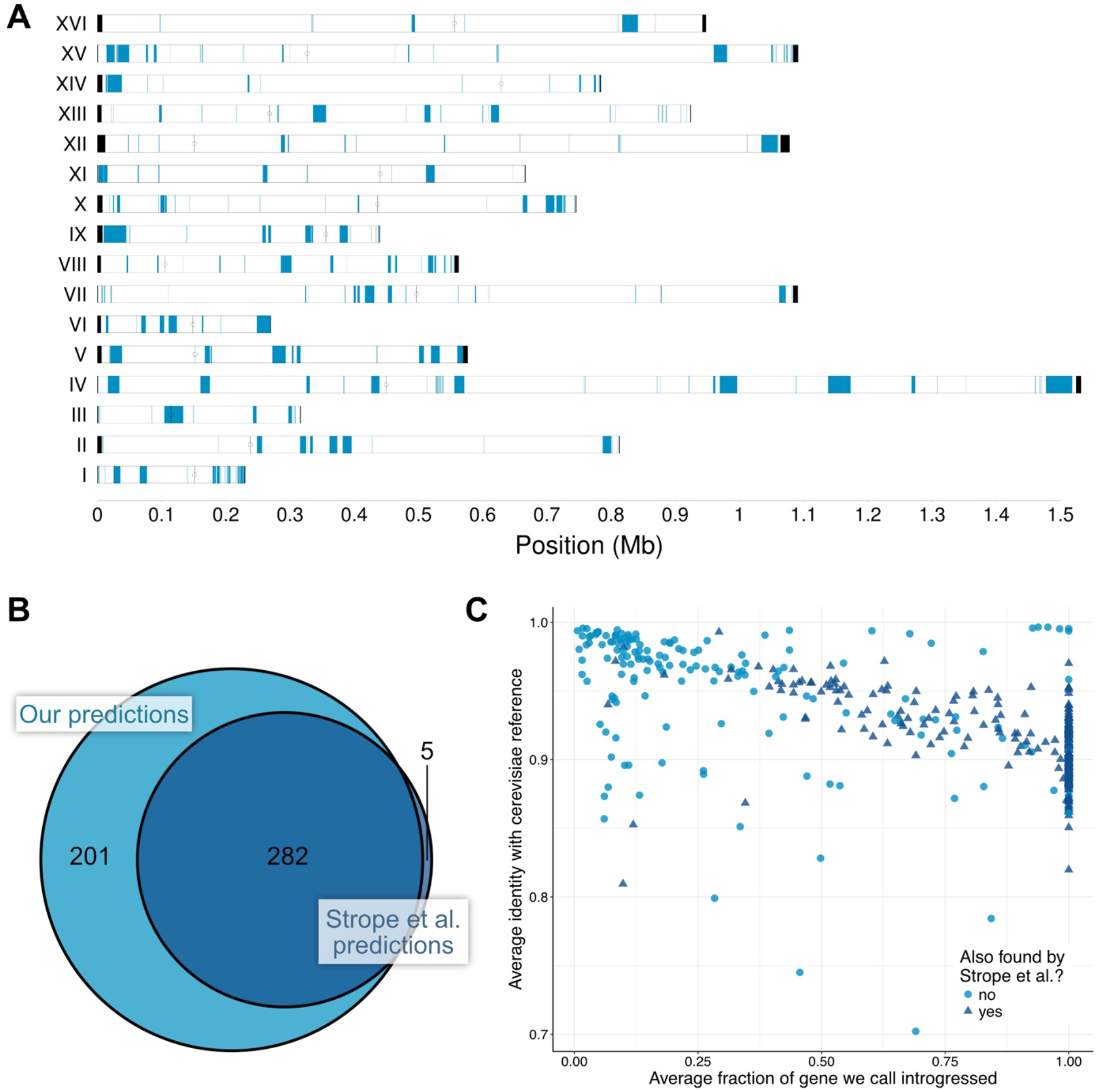
The introgressed regions and genes we identify, compared to previously identified genes. (*A*) The distribution of introgression across the entire genome. Regions that we identify as introgressed in one or more strains are colored blue. Telomeres are colored black, and centromeres are marked with a black line and circle. (*B*) The introgressed regions we identify overlap a total of 483 genes, 282 of which were previously identified as introgressed *(Strope et al. 2015)*. We fail to find five previously-identified genes because of translocations. (*C*) The genes we newly identified as introgressed tend to have only a fraction of their sequence introgressed, resulting in greater identity with the *cerevisiae* reference across the whole gene sequence.

We were concerned that some of the genes we identified might have been subject to paralogous gene conversion rather than introgression. To evaluate whether this was the case, we examined the 104 genes we identified that had a paralog. We used BLAST (Altschul et al. 1990) to compare the introgressed portion of each gene to (1) the gene from the *S. cerevisiae* reference, (2) the gene from the *S. paradoxus* reference, (3) the paralog from the *S. cerevisiae* reference, and (4) the paralog from the *S. paradoxus* reference. We then examined the genes for which the best hit was in the *S. cerevisiae* paralog (rather than the *S. paradoxus* gene, as we would expect for introgression). In the case of four genes (*RPL8B, SSB2, SSF1*, and *SSF2*), paralogous gene conversion appeared to be a more likely explanation than introgression. Thus, this phenomenon does appear to result in some false positives in our predictions, but is not likely to be responsible for a substantial fraction of the paralogs we identify as introgressed.

### Introgression in specific strains

The amount of the genome we predict to be introgressed ranges from 0.03% to 4.4% across all the strains (Figure 4a). As noted previously (Strope et al. 2015), strains YJM1252, YJM248, and YJM1078 appear to have far more introgression than the other strains. Much of this introgression is shared (Figure 4b) and is comprised of longer regions (Figure 4c), suggesting that an ancestor of these three strains hybridized with *S. paradoxus* relatively recently.

**Figure 4.**
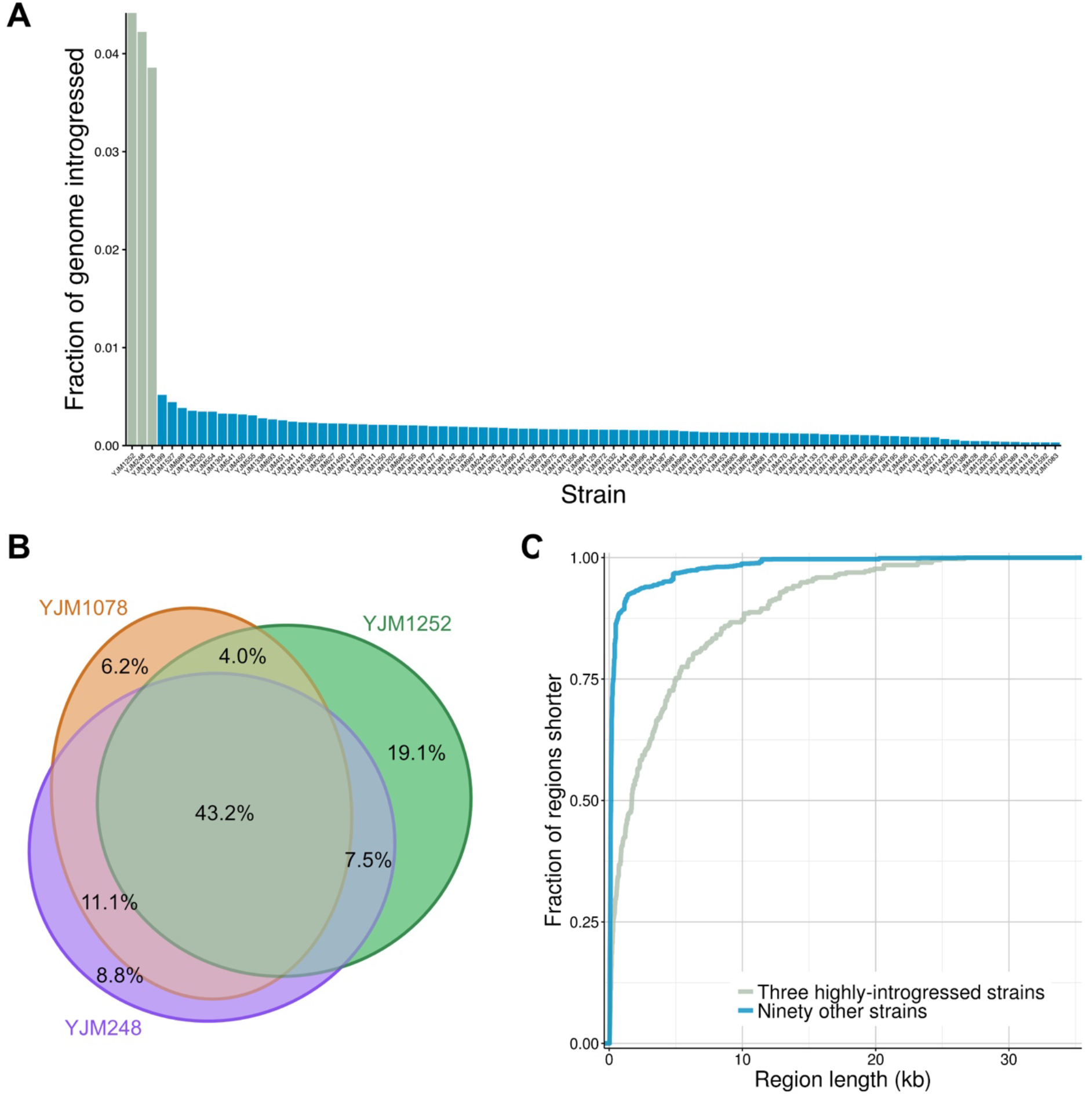
Introgression across all strains, and in three highly–introgressed strains. (*A*) We predict less than half a percent of most genomes to be introgressed from *S. paradoxus*, but three strains have a much higher level of introgression. (*B*) Much of this introgression is shared among all three strains, and about two-thirds is shared between at least two of the strains. (*C*) Most of the introgressed regions we find are short (less than 150bp), but these three highly-introgressed strains have many longer introgressed regions, including a quarter of the regions that are greater than 5kb in length.

Most of the introgressed genes we identify are introgressed in only a few strains, but some are more widespread (Figure 5). The most frequent number of strains for an introgressed gene is three because of the large amount of introgression shared between the three strains mentioned above; however, there are also many examples of gene introgressions shared among other sets of strains. For example, the same small part of the gene *SIR4*—which is involved in the assembly of silent chromatin domains—is introgressed in YJM1199, YJM1202, and YJM1304. A 603bp introgressed region of this gene has introduced 142 polymorphisms and one 3bp insertion into these three strains, resulting in 82 amino acid changes and one insertion.

**Figure 5.**
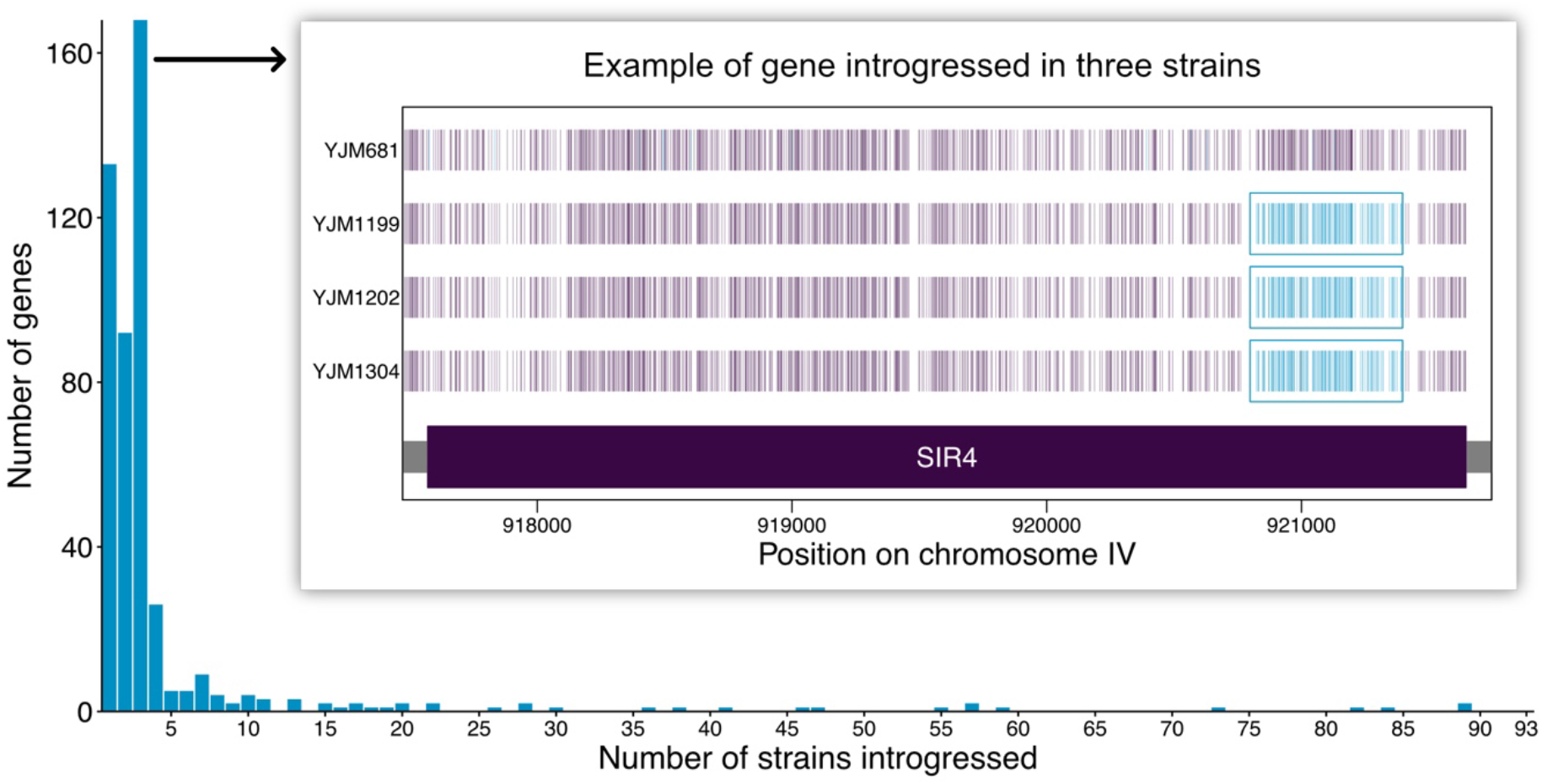
The number of strains each introgressed gene is found to be introgressed in. Inset shows an example of one gene, SIR4, that is partially introgressed in three strains. A strain without introgression in this gene is shown for comparison. For each strain, vertical lines indicate alignment columns at which the strain matches the *S. cerevisiae* reference but not the *S. paradoxus* reference (purple), the *S. paradoxus* reference but not the *S. cerevisiae* reference (blue), or neither reference (gray). Alignment columns containing gaps or ambiguous sequence calls are not shown.

### Impact of introgression on global polymorphism and phylogenetic inference

Although we only predict a small fraction of most *S. cerevisiae* genomes to be introgressed from *S. paradoxus*, the level of divergence between the two species suggests that this small amount of sequence could account for a relatively larger amount of the nucleotide diversity present in *S. cerevisiae*. We calculated nucleotide diversity among the 93 strains across the entire genome to be 0.81%. We separately calculated nucleotide diversity excluding sites predicted to be introgressed to be 0.76%, indicating that 6.6% of the nucleotide diversity is contributed by these putatively introgressed regions (Figure 6a). Repeating the same analysis only in coding regions, we find that slightly less (5.5%) of the nucleotide diversity is contributed by the introgressed regions, consistent with the hypothesis of purifying selection acting more strongly on protein versus non-coding regions. Furthermore, there is marked heterogeneity in the contribution of introgressed sequences to nucleotide diversity across chromosomes (Figure 6a).

**Figure 6.**
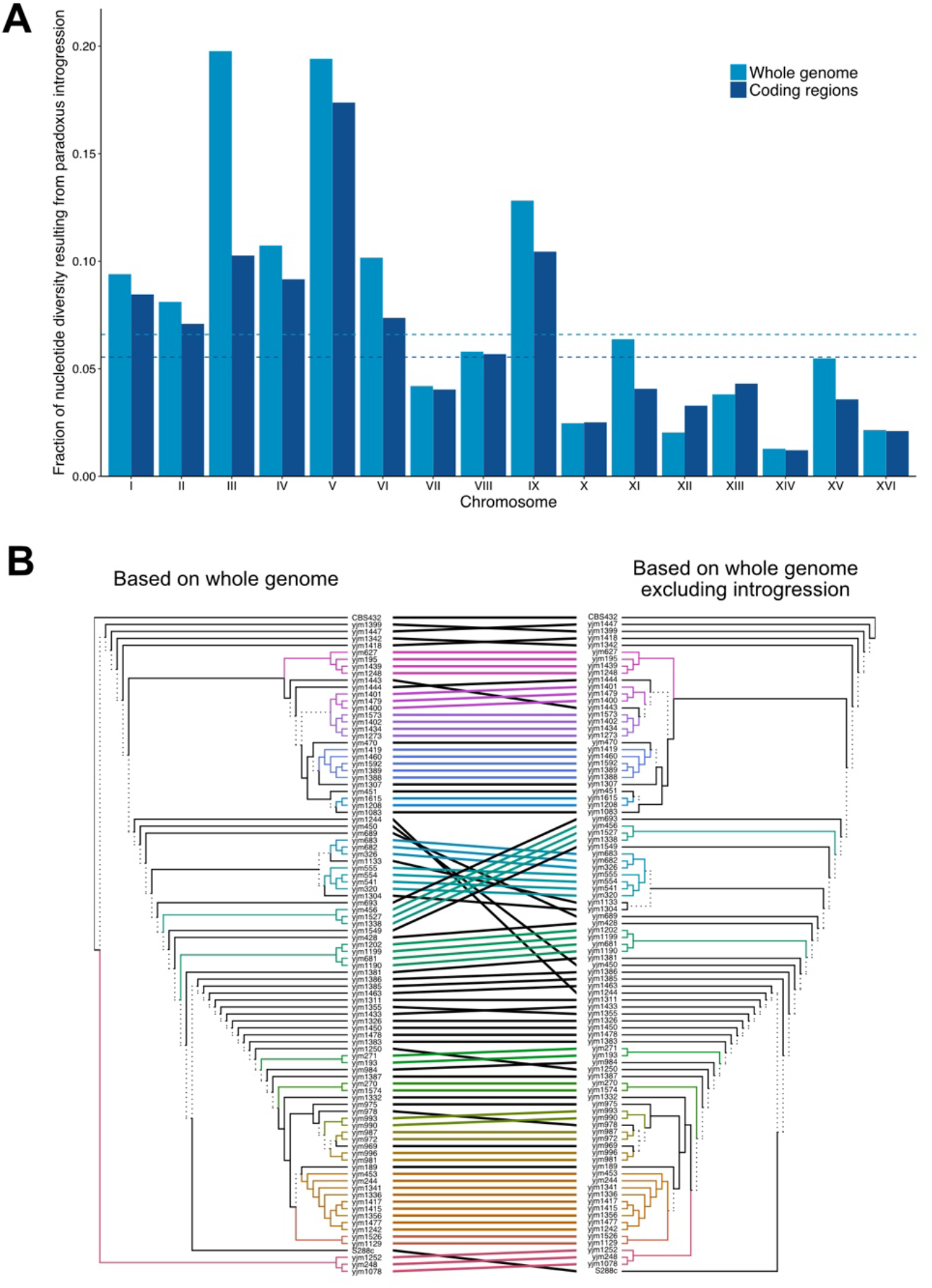
The impact of introgression on overall diversity and phylogenetic relationships within *S. cerevisiae*. (*A*) The fraction of nucleotide diversity contributed by introgressed regions on each chromosome, for all sites and all coding sites. Genome-wide averages are shown with dotted lines. (*B*) Tanglegram of phylogenies constructed from sites across the entire genome and from only non-introgressed regions. The whole-genome phylogeny is a consensus tree of 50 jackknife replicates generated by removing a number of random sites equal to the number of introgressed sites removed for the non-introgressed tree.

Given the relatively large effect of these introgressed sequences on nucleotide diversity, we next tested whether failing to account for introgression would influence inferences of the evolutionary history of these strains. We found that excluding introgressed regions resulted in a substantially different phylogeny than one constructed using all sites (Figure 6b). Unsurprisingly, removing introgressed sites moves the three highly–introgressed strains YJM1252, YJM248, and YJM1078 to be more closely related to other *S. cerevisiae* strains, but several other strains also move to different clades in the phylogeny for less obvious reasons.

We were also interested in whether introgressions cluster together on a phylogeny of these species, which would indicate they may have come from common hybridization events. We looked at all genes that overlapped an introgressed region in at least one strain, and clustered the genes based on their phylogenetic distribution, using a tree constructed from sites without introgression (Supplemental Figure 3). If most introgressions are due to just a few hybridization events, we would expect to see distinct clusters falling within clades. There are some examples of such patterns, and further analysis of specific shared introgressions may allow for the inference of when hybridizations are likely to have occured.

## Discussion

We have implemented a simple, flexible, and powerful method for identifying introgression among the *Saccharomyces* yeasts. By applying this method to look for *S. paradoxus* introgression in a set of 93 geographically and ecologically diverse *S. cerevisiae* strains, we have found introgression to be pervasive but highly variable across strains.

Although we have identified some amount of introgression in all of these strains, it is important to recognize that our inferences are dependent on our choice of reference genomes. In particular, we are unlikely to identify introgressions in *S. cerevisiae* strains that are also present in the *S. cerevisiae* reference we are using. Using a different *S. cerevisiae* reference genome may allow us to identify additional introgressions; however, sequence identity–based approaches like ours will always struggle to identify introgressions that are relatively old or that fall within highly– conserved regions of the genome. Conversely, our set of introgressed regions likely also contains some false positives. We have filtered out many of these, and the ones that remain are on the whole more likely to be due to poorly–aligned regions than to incomplete lineage sorting or paralogous gene conversion.

Our analysis of introgression has so far only concerned the presence of *S. paradoxus* sequence in *S. cerevisiae* genomes, but we would expect to find evidence of introgression in the opposite direction, as well. Furthermore, there is a substantial amount of population structure within *S. paradoxus*, with divergences between its different populations ranging from ~1-3% (Leducq et al. 2014). Our model can be easily extended to include additional *S. paradoxus* reference genomes or even references for other *Saccharomyces* species; such an expanded analysis could allow us to identify more introgression, and perhaps to gain a clearer picture of when and where hybridization occurs.

Although we have mainly focused in this study on general patterns of introgression among the strains we analyzed, our method may also allow for the identification of specific examples of adaptive introgression that could be further characterized experimentally. Overall, we hope this approach may yield further insights into the evolution of *Saccharomyces* yeasts, and specifically the role that hybridization has played and continues to play in their adaptation to new environments.

## Methods

### Analytical theory to estimate probability of incomplete lineage sorting

The probability of incomplete lineage sorting (ILS) is given by one minus the probability that a given gene tree and species tree are concordant. The formula for this probability can be derived using coalescent theory, and depends on the number of samples for each species and the scaled divergence time between the species (Rosenberg 2002). The scaled divergence time is given by *T* = *t/N*, where *t* is the divergence time in generations, and *N* is the effective population size of each species. We are specifically interested in monophyletic concordance, the scenario in which both the gene tree and species tree are monophyletic. The probability of monophyletic concordance can be calculated as

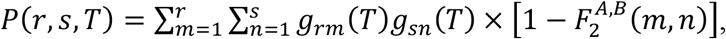

where *g*_*ij*_ (*T*) is the probability that *i* lineages derive from *j* lineages that existed *T* coalescent time units in the past, and is given by

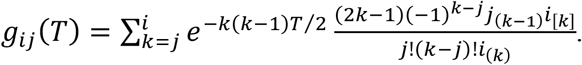

The quantity 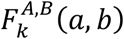 is the probability that an interspecific coalescence occurs in coalescing *a* and *b* lineages from species *A* and *B* respectively to *k* total lineages:

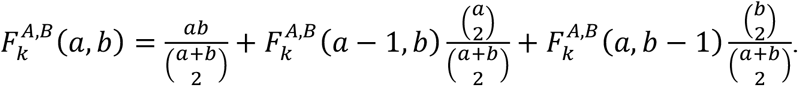

To estimate the probability of ILS when looking at *S. paradoxus* and *S. cerevisiae* sequences, we assume one individual from the former species and 94 individuals from the latter, to correspond to the numbers of genomes we examine later. We estimate the time, *T*, from coalescent simulations (described in more detail in Simulated Sequences) that produce the expected approximate sequence identity between the species of 86%. These simulations require as parameters the effective population size and mutation rate. The effective population size of *S. paradoxus* has been estimated as 8.6×10^6^ for the European clade and 7.2×10^6^ for the Far Eastern clade, and these estimates are in turn based on estimates of the mutation rate and inbreeding coefficient (Tsai et al. 2008). We set the effective population sizes for *S. cerevisiae* and *S. paradoxus* as 8×10^6^ individuals each. Using a mutation rate of 1.84×10^−10^ (Fay and Benavides 2005), we estimated the time of divergence as 3.75×10^8^ generations.

### Hidden Markov model

The hidden Markov model has one state for each reference species as well as one unknown state. The model was used to assign a state to individual alignment columns in each three-way whole chromosome alignment. Only alignment columns that were polymorphic and contained no gaps were considered. The initial, emission, and transition probabilities were set by a combination of (1) initial estimates of the number and sizes of introgressed regions, and (2) the number of alignment columns in which the test strain matched each of the reference strains. These probabilities were updated by Baum-Welch training until the log likelihood decreased by less than 0.1% between iterations.

In analyzing simulated sequences, alignment columns with a high posterior probability (above a given threshold) of being in the paradoxus (introgressed) or unknown state were assigned those respective states, and were assigned the cerevisiae (non-introgressed) state otherwise. In analyzing actual chromosomes, the Viterbi algorithm was used to find the most likely state sequence. In both cases, introgressed regions were defined as continuous blocks of alignment columns assigned to the introgressed state, including any intervening alignment columns that were monomorphic or contained gaps. All regions are indexed relative to the *S. cerevisiae* reference genome.

### Simulated Sequences

Sequences were simulated using ms (Hudson 2002, https://uchicago.app.box.com/s/l3e5uf13tikfjm7e1il1eujitlsjdx13). Two samples were simulated for S. *cerevisiae*, one for S. *paradoxus*, and one for an outgroup. The effective population size was set to 8×10^6^ for all populations. Migration rate, expressed as the fraction of the *S. cerevisiae* population made up of *S. paradoxus* individuals in each generation, was set to either zero or a value ranging from 5×10^−10^ to 5×10^−8^, and was set to only occur in the most recent 5% of time since the species diverged. The recombination rate was set to 7.425×10^−6^, which was calculated from the formula 1 + 6.1 crossovers/Mb (Mancera et al. 2008), using the average *S. cerevisiae* chromosome size. The mutation parameter, Θ = 4N0µ, was set using a mutation rate of µ=1.84×10^−10^ (Fay and Benavides 2005). Simulated sequences were 100,000 bp in length. An example ms command with a migration rate of 5×10^−10^ is:

`ms 4 100 -t 5.888 -r 0.09494496 100000 -p 8 -I 3 2 1 1 -m 1 2 0.32 -m 1 3 0.0 -em 0.5859375 1 2 0 -em 11.71875 1 3 0 –ej 11.71875 2 1 -ej 35.15625 3 1 -T`

### PhyloNet-HMM

PhyloNet-HMM (Liu et al. 2014) was run with trees of three species (representing *S. cerevisiae, S. paradoxus*, and an outgroup), for a total of six possible states for each alignment column. Parameter input files are provided in Supplemental File 1. Sequences were generated from the ms simulation output using seq-gen (https://github.com/rambaut/Seq-Gen/releases/tag/1.3.4).

### Strain genomes, annotations, and alignments

The reference *S. cerevisiae* S288c and *S. paradoxus* CBS432 genomes were downloaded from SGD (https://downloads.yeastgenome.org/sequence/). The 93 non-reference strains were downloaded from GenBank (https://www.ncbi.nlm.nih.gov/genbank/) based on the table of accession numbers provided by Strope et al., 2015. The lists of all verified ORFs and all paralogs in *S. cerevisiae* S288c were downloaded from SGD using YeastMine. Three-way alignments between the *S. cerevisiae* and *S. paradoxus* reference genomes and each *S. cerevisiae* test strain were performed on each chromosome separately using MAFFT (Katoh and Standley 2013) with default settings. The two reference genomes were also aligned separately. Changing the ep parameter from 0.0 to 0.321 did not result in substantially shorter alignments.

### Filtering

Low-complexity regions of individual genomes were masked using dustmasker from BLAST version 2.7.1 (ftp://ftp.ncbi.nlm.nih.gov/blast/executables/blast+/2.7.1/). Introgressed regions predicted by the HMM were included in final analysis if and only if they met all of the following criteria:

- the fraction of alignment columns containing at least one gap or at least one masked base did not exceed 0.5,
- the number of sites at which the test strain matched the *S. paradoxus* reference but not the *S. cerevisiae* reference was at least seven, and
- the divergence between the test strain and the *S. cerevisiae* reference (calculated on all alignment columns without gaps or masked bases) was less than 0.3.

### Phylogenies

A FASTA file was generated with a position for every site in the reference *S. cerevisiae* S288c genome. The corresponding nucleotides for the other 93 *S. cerevisiae* strains were extracted from the three-way alignments, and for the *S. paradoxus* CBS432 reference from a two-way alignment. All alignment columns with gaps were removed, leaving a total of 6,033,510 sites. Then, all 376,536 of these sites that overlapped an introgressed region in any strain were removed, leaving a total of 5,656,975 sites. A phylogeny was constructed from these non-introgressed, non-gapped sites.

Phylogenies were constructed using PHYLIP (http://evolution.genetics.washington.edu/phylip/) dnadist and neighbor, using the UPGMA algorithm. In addition, a phylogeny was constructed in the same way from a dataset generated by removing a random set of sites equal in size to the set of introgressed sites; this sampling was repeated 50 times using PHYLIP seqboot, and a consensus tree was generated using PHYLIP consense. The tanglegram was plotted using the R package dendextend (Galili 2015).

## Data access

The implementation of our hidden Markov model and downstream analyses are available for download on GitHub (https://github.com/hyperboliccake/introgression).

## Notes

### Competing Interest Statement

The authors have declared no competing interest.

