## Supplementary figures and images for "The genomic landscape of *Saccharomyces paradoxus* introgression in geographically diverse *Saccharomyces cerevisiae* strains"

### Supplemental Figure S1

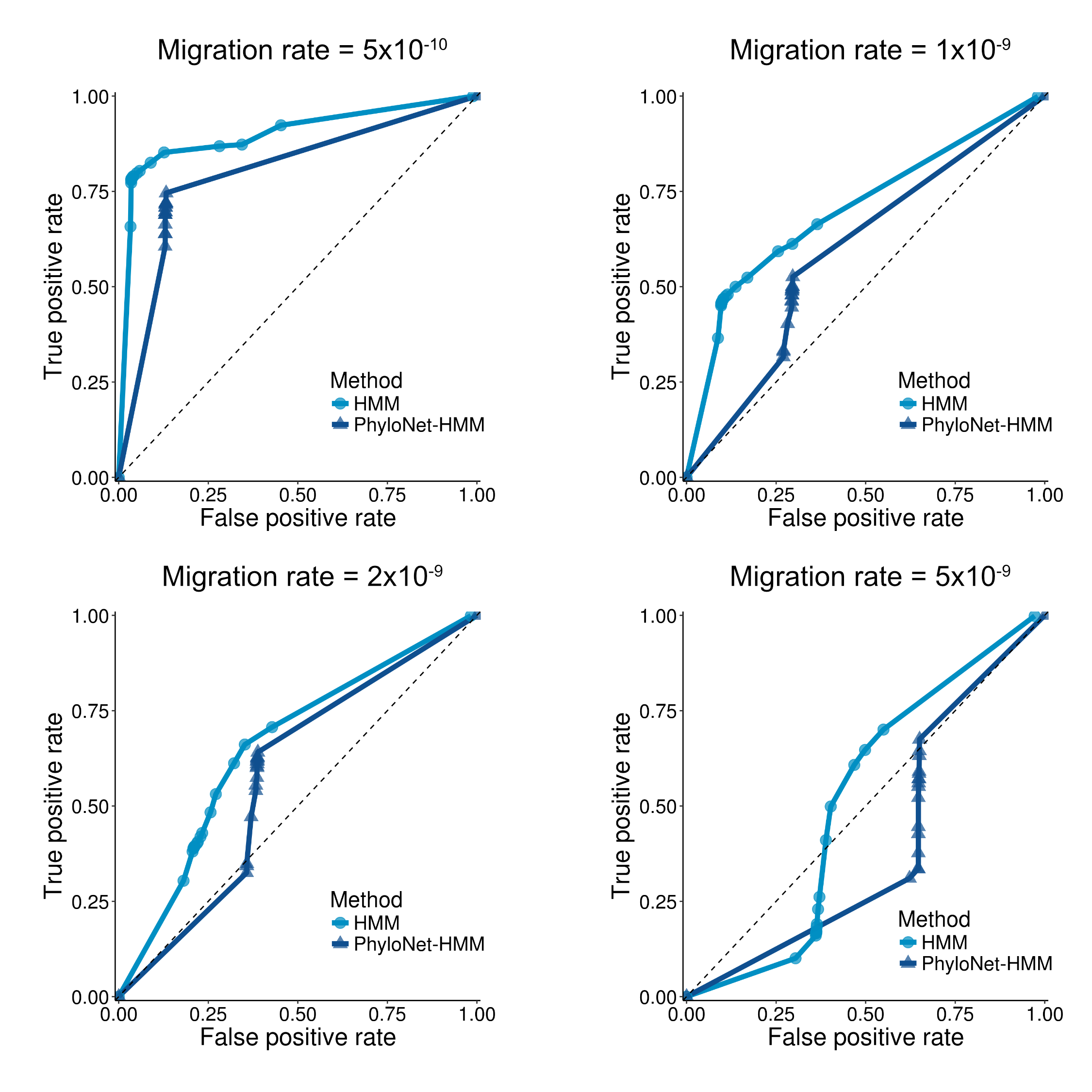

### Supplemental Figure S2

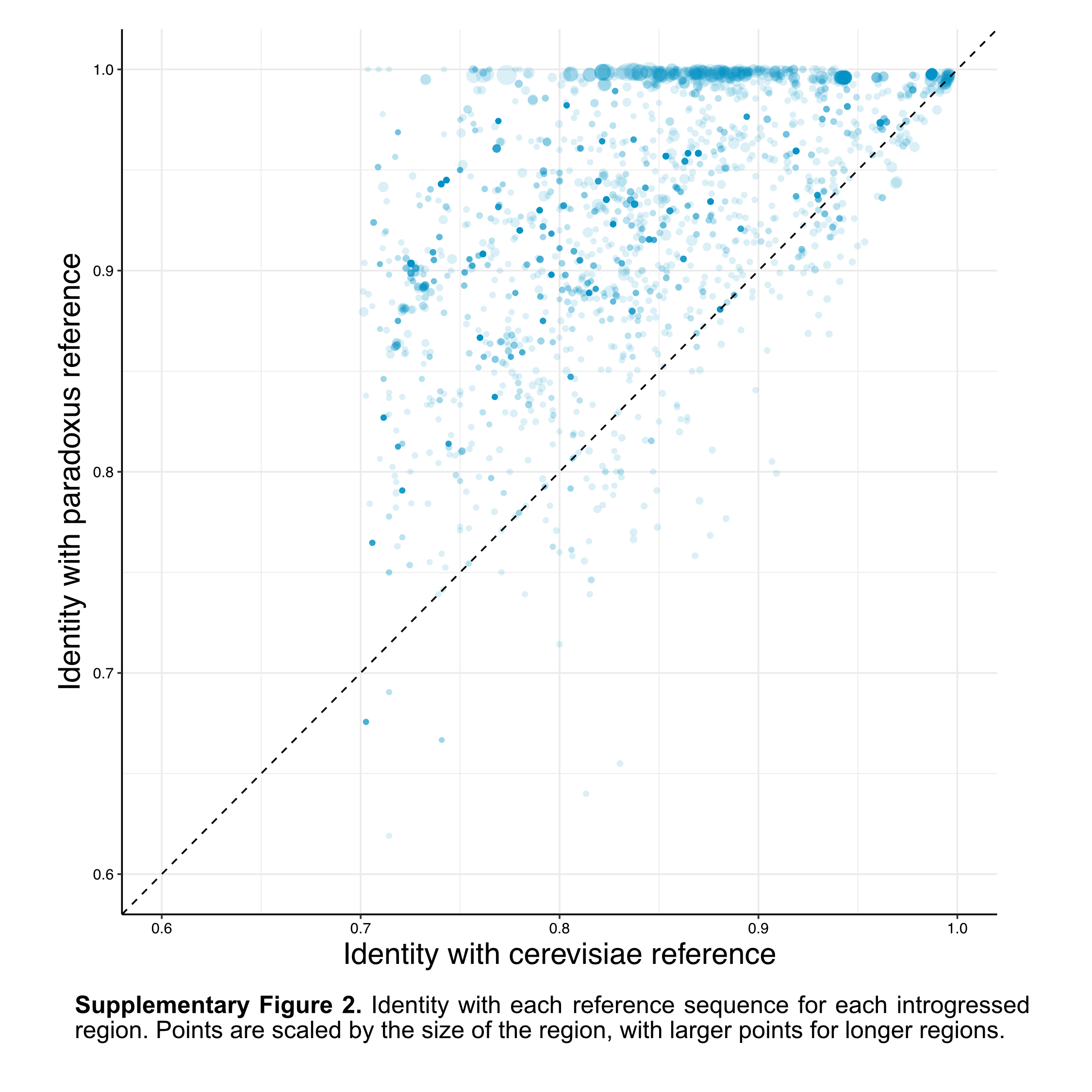

### Supplemental Figure S3

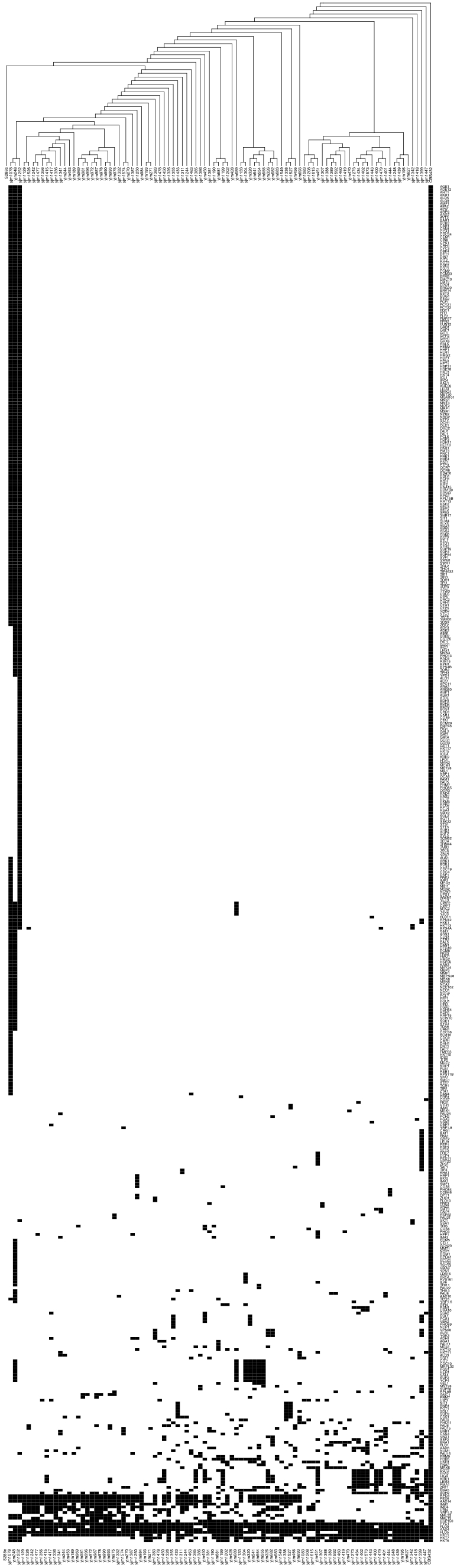
